# Chromosome interactome inferred from mitosis-G1 transition

**DOI:** 10.1101/084608

**Authors:** Y.A. Eidelman, S.V. Slanina, A.V. Aleshchenko, S.G. Andreev

## Abstract

The progress in experimental techniques aimed at 3D genome study is yet to bring about revelation of basic principles of genome folding. Chromosome conformation capture Hi-C technologies provide genome wide mapping of genomic loci interactions but spatial organization of chromosomes remains unknown. Here, we develop a polymer modeling approach to generate the ensemble of 3D chromosome conformations for mapping genetic loci contacts and the positions of megabase chromosomal domains in interphase chromosome at different time of mitosis-interphase transition. We demonstrate that (*) whole chromosome contact map (interactome) generated for mouse chromosome 18 structure and (**) contact patterns, observed soon after mitotic decondensation and remaining similar during G1, correlate well with the experimental Hi-C contact data. The results suggest that contact map formation and spatial compartmentalization of an interphase chromosome are driven by interactions between different types of domains during formation of globular chromosome state at the end of mitotis-G1 transition.

## Introduction

Whole genome chromosome conformation capture (Hi-C) based experiments provide unique information about frequencies of contacts of any genetic loci within and between chromosomes in a cell. In general, the method represents a physico-chemical approach which uses chromatin fragmentation and proximity ligation and gives a snapshot of crosslinked chromosomal contacts in the population of fixed cells in the form of contact maps (Lieberman-Aiden et al., 2009, Dixon et al., 2012, Rao et al., 2014, Schmitt A.D. et al 2016).

However Hi-C technology has some inherent drawbacks. One of them is inability to provide direct information about 3D structures (Lesne et al., 2014, Ay and Noble, 2015). To infer 3D chromosome structures from Hi-C data complementary computational approaches are extensively developed, see reviews (Ay and Noble, 2015, Imakaev et al., 2015, Rosa and Zimmer, 2014, Schmitt A.D. et al 2016). They are divided into two main groups. One group reconstructs conformations of chromosomes basing on the phenomenological relationship between contact frequency and spatial distance between loci (Lesne et al., 2014, Ay and Noble, 2015, Schmitt A.D. et al 2016). This type of methods is limited to prediction of single “average” geometrical model of structure and unable to generate long-range chromosome interaction maps observed by Hi-C. Another group of methods is based on statistical polymer physics and is able to predict ensembles of 3D conformations and contact maps, which are sensitive to principles of organization. Therefore polymer modeling of chromosomes serves as an effective tool to deduce principles of genome organization from quantitative processing of Hi-C information (Imakaev et al., 2015, Schmitt A.D. et al 2016). Studies aimed at polymer modeling of interphase chromosomes for Hi-C analysis use condensation algorithms (Andreev and Edel’man, 1999, Rosa and Zimmer, 2014, Zhang and Wolynes, 2015, Brackley et al., 2016, Pierro et al., 2016, Chiariello et al., 2016). Simplistically, the condensation conditions for long polymer chromatin chain due to bridging proteins (Brackley et al., 2016, Chiariello et al., 2016) or attracting potentials (Andreev and Edel’man, 1999, Zhang and Wolynes, 2015, Pierro et al., 2016) are applied and large scale transition from extended chromatin fiber, as rod or coil, to compact state, associated with interphase chromosome is simulated. These assumptions contradict what happens in living cells where chromatin/chromosome large scale condensation is not responsible for establishment of G1 chromosome organization. It is common knowledge that interphase chromosome territories are formed as a result of decondensation of mitotic chromosomes during transition to G1. Therefore inferring chromosome structure on the basis of polymer physics faces the problem whether polymer methods can be applied to the cell.

To overcome methodological issues we present the new modeling approach aimed at predicting interphase chromosome architecture by means of evaluation of chromosome dynamics during mitosis-G1 transition. To explore complexity of chromosome folding we map genetic loci contacts and predict positions of megabase chromosomal domains in interphase chromosome at different time of mitosis-interphase transition by Monte-Carlo simulations. Generated chromosomal contact frequencies for mouse chromosome 18 in interphase state (interactome) are demonstrated to be in good consistency with the experimental Hi-C data at megabase resolution. We conclude that chromosomal domains are organized into four types of compartments which are different in size and 3D configuration. To examine validity of condensation-based polymer methods we reproduce the condensation scenario, predict and compare parameters of chromosome structure for both approaches. Taken together, our results support a mechanism of chromosome folding underlying contact patterns formation via interactions between different types of domains during establishment of globular chromosome state at the end of mitotis-G1 transition. This mechanism consistently reproduces a complex chess-like pattern and early G1 establishment of long-range contacts as well as spatial compartmentalization of chromosome under different conditions.

## Results

Using Monte Carlo (MC) simulations, we investigate the establishment of interphase chromosome organization in the course of mitosis-G1 transition. At t=0 decondensation begins, the chromosome is extended to the chain of subunits representing megabase-sized domains confined by external field (see Materials and methods). These structural chromosome subunits can be interpreted as topological domains, TDs (Dixon et al., 2012), topologically associated domains, TADs (Nora et al., 2012) or superdomains, SDs (Eidelman and Andreev, 2002). According to the presented model, domains interact with each other in a chromosome via interactions between non-neighboring subunits. They are considered in statistical polymer physics as volume interactions (Lifshitz, 1969) and hereafter we use this term. These factors shape 3D conformations during mitotic chromosome decondensation and in the interphase.

The simulated dynamics of mouse chromosome 18 structure during mitosis-G1 transition is shown in Figure 1a. Dynamics of chromosome gyration radius during mitosis-G1 transition, Figure 1b, curve 1, indicate that quick growth gives way to slow decline. At the slow phase, chromosome conformation rearrangements accompanied by subcondensation take place. At intermediate/late time scale (50,000 steps and more) the decline is quite small. To determine the type of interphase chromosome structure we calculated the radial dependence of density of chromosomal subunits at 50,000 and 500,000 MC steps, Figure 1c. The results suggest that the interphase chromosome is a globule with the dense core and the rather loose fringe.

**Figure 1.**
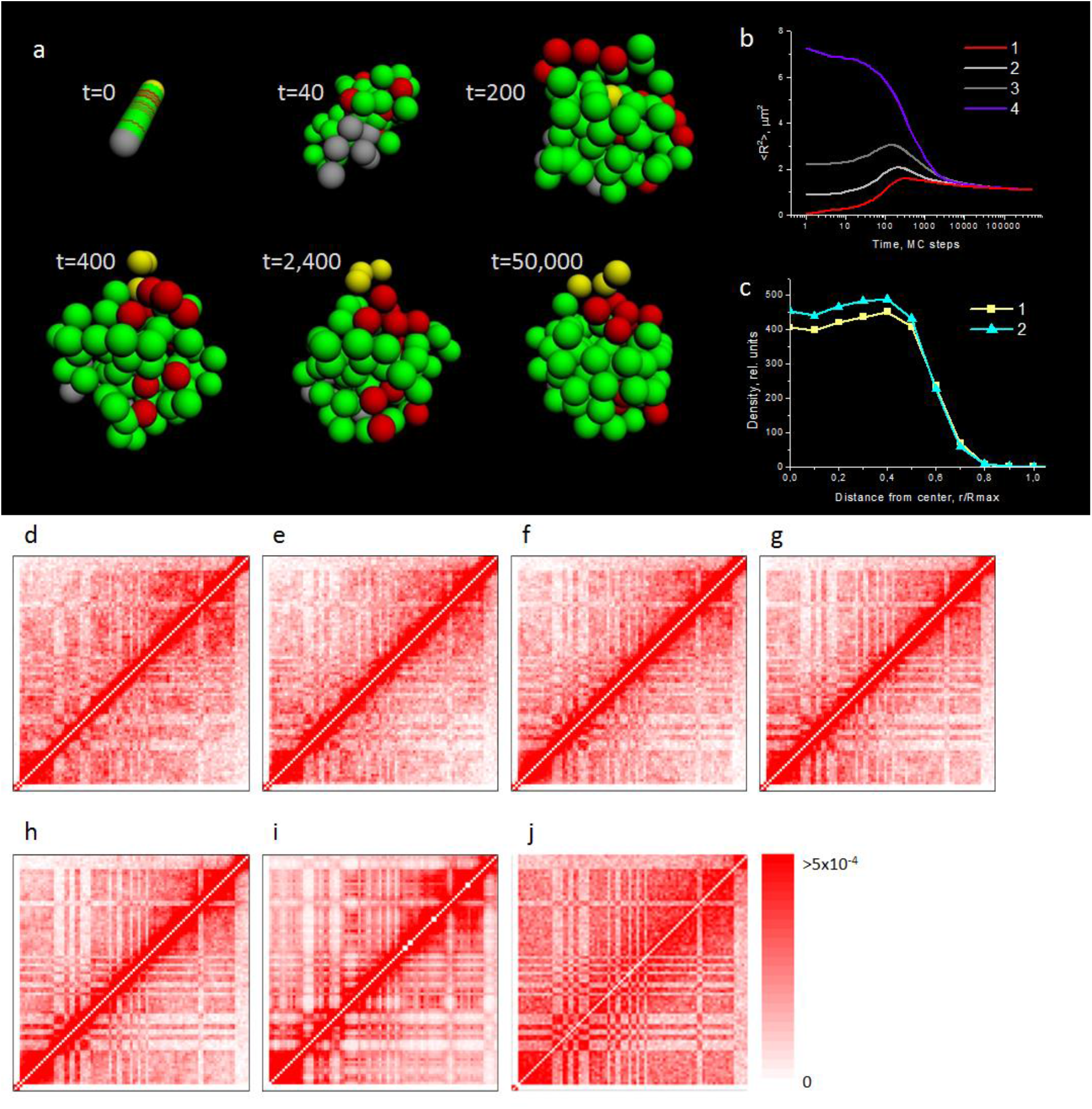
Globular structure of mouse chromosome 18 established as a result of mitosis-G1 transition. **a** Evolution of the typical conformation in the course of mitosis-G1 transition. The times shown are in MC steps. Colors: yellow – centromeric subunits; green – subunits type 1; red – type 2; grey – type 3. t=0 corresponds to the mitotic-like conformation, t=50,000 corresponds to the interphase conformation. **b** squared gyration radius dynamics during mitosis-G1 transition. 1-3 – decondensation from mitotic-like conformation of different density. 1 – distance between neighboring subunits 5 nm (see t=0 in (a)) 2 – 25 nm; 3 – 40 nm. 4 – condensation from polymer coil. **c** radial density distribution in the interphase state. 1 – t=50,000 MC steps. 2 – t=500,000 MC steps. **d**-**j** contact maps for mouse chromosome 18. **d**-**h** map simulations for different times. **d** t=2,400 MC steps. **e** t=5,000 MC steps. **f** t=10,000 MC steps. **g** t=50,000 MC steps. **h** t=500,000 MC steps. **i** experimental map (Rao et al., 2014) for chromosome 18, ch12lx mouse cells. Pearson correlation between **h** and **i** R=0.897. **j** maps for “broken chain” model. Upper triangle: t=50,000 MC steps, lower triangle: t=500,000 MC steps. All maps are drawn with 1 Mbp resolution. Contacts with |*i-j*|=0, i.e. within 1 Mbp, are not counted (white diagonal lines). In all cases 500 structures are simulated

To elucidate how dynamics of gyration radius depends on scenario of mitosis-G1 transition we performed calculations for other models of mitotic chromosome decondensation (see Materials and methods), with less compact initial state, Figure 1b, curves 2 and 3. Time dependence of gyration radius is changed, initial sharp increase of gyration radius becomes less expressed. The results suggest that early appearance of contact pattern may depend on mechanisms of mitotic decondensation.

To assess time dependence of contact map establishment during G1 we calculate contact maps at different times after mitotic decondensation: at early (2,400 and 5,000 MC steps), intermediate (10,000 and 50,000 MC steps) and late times (500,000 MC steps) (Figure 1d-h). We associate these points with different times in G1, from early to late subphases. Strictly speaking, the boundary of G1 is not determined here and onset of S phase is not considered in the model. Therefore we use the simplified, slightly arbitrary, classification of G1 subphases. The term “early” G1 is introduced here by the simple morphological criterion as disappearance of mitotic chromosome conformation and appearance of interphase-like conformation. This criterion does not depend on absolute scale of MC steps vs real time. Thus an early G1 map is a map for chromosome after decondensation and the chromosome shape is a globule morphologically similar to that at late times in G1 (Figure 1a). Map calculations for different times, Figure 1d-h, show that the chess-like contact pattern is less definitely manifested at 2,400 MC steps and becomes clearer at 5,000 and 10,000 MC steps. But positions of characteristic dark/light blocks at early time remain unchanged at intermediate and late times. After 50,000 steps the map visually ceases to change. Further calculations at 500,000 (Figure 1h) and at 10,000,000 steps (not shown) result in the maps practically indistinguishable from that at 50,000 steps. These data reveal that early G1 contact maps (at 2,400 and 5,000 MC steps) are highly correlated with the later time contact maps (R=0.938 for 2,400 vs 50,000, R=0.923 for 2,400 vs 500,000, R=0.963 for 5,000 vs 50,000, R=0.952 for 5,000 vs 500,000 MC steps).

The group of Hi-C data (Naumova et al., 2013) was obtained with the use of cell synchronization, that allows to distinguish between early and late G1. The conclusion (Naumova et al., 2013) is that contact maps are highly correlated in early vs mid G1 vs early S. Our numerical data agree with this finding.

Quality of the contact map in G1 generated by the polymer modeling is demonstrated by comparison with the experimental map derived from Hi-C data (Rao et al., 2014), Figure 1i. The contact map predicted for 50,000 (as well as for 500,000 and 10,000,000) MC steps is highly correlated with the experimentally derived Hi-C map, Pearson correlation coefficient R=0.897. Thus main patterns of the experimental map^3^ are well reproduced by polymer modeling of interphase chromosome 18, established following mitosis-G1 transition.

To explore origin of chess-like pattern of interactions observed in the experimental map (Figure 1i), we computed the contact maps for the models of a homopolymer globule and of a heteropolymer globule decondensed to coiled state, Supplementary Figure 1a,b. Both models provide no chess-like pattern of contacts. Thus the globular state of the heteropolymer (block-copolymer) chromosome is the necessary condition for the chess-like structure of the contact map.

Next, we examined the impact of external field on map compartmentalization. Elimination of external field results in partial loss of long-range contacts, but the chess-like pattern remains, Supplementary Figure 2a,b. Thus external field, or confined chromosomal environment, serves as an important factor for reproducing the complex pattern of contacts.

**Figure 2.**
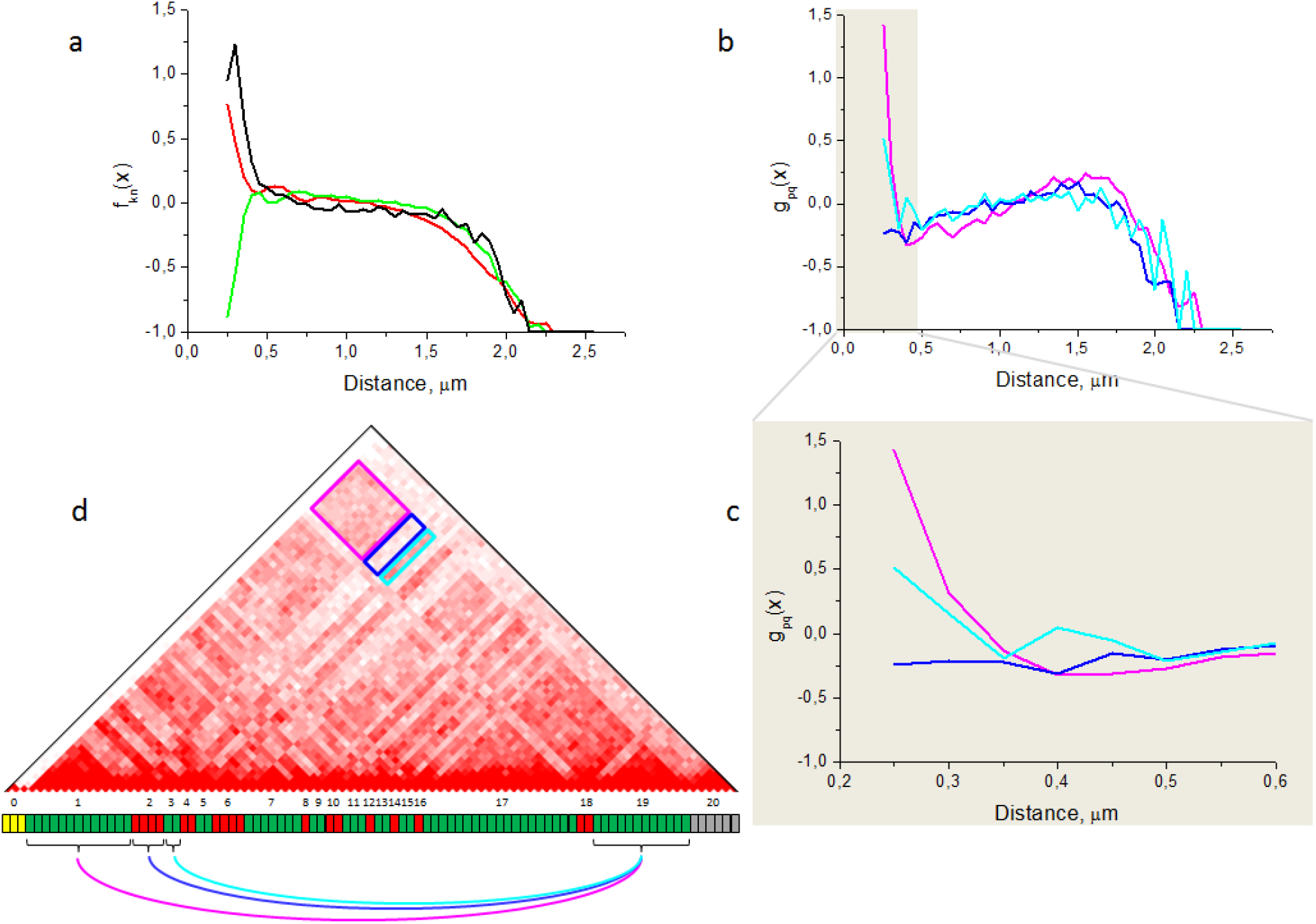
Spatial compartmentalization of chromosome 18. **a** relative difference function f_kn_(x) between T_kn_(x) and T(x). Red curve: k=1, n=1; black: k=2, n=2; green: k=1, n=2. Functions T_kn_(x) and T(x) are presented in Supplementary Figure 4. **b** relative difference function g_pq_(x) between F_pq_(x) and T_long_(x). Magenta: p=1, q=19 (i.e. compartment 1 vs compartment 19). Blue: p=2, q=19. Cyan: p=3, q=19. **c** same as **b** but only the short-distance region is shown. **d** schematic representation of compartment interactions. The contact map is the same as in Figure 1g, with regions corresponding to compartment pairs 1-19, 2-19 and 3-19 marked by colored frames. The chromosome ideogram below shows compartment layout, with compartment numbers indicated straight above (the centromere is marked as compartment 0). Colored arcs indicate interactions between compartment pairs 1-19, 2-19 and 3-19. Compartment 1: 3-16 Mbp; compartment 2: 16-20 Mbp; compartment 3: 20-22 Mbp; compartment 19: 73-85 Mbp

To further explore contribution of different factors to contact map compartmentalization we introduce the “broken chain” approach. This means that the polymer chain is artificially cut between subunits and the chromosome is represented by a gas of independent subunits. As previously, there are volume interactions and external field confinement, but now there is no spatial correlations of megabase subunits along the chromosome, or no linear memory in terms of theory of globular state (Lifshitz, 1969). The contact maps calculated for this artificial system at different times, 50,000 and 500,000 MC steps, are shown in Figure 1j. It is clearly seen that map compartmentalization is mainly preserved although some differences, mainly in short-distance contacts, are also observed with respect to intact unbroken polymer chain (Figure 1g,h). The weak difference indicates the minor contribution of subunit connectivity, or linear memory to the complex pattern of contacts for mouse chromosome 18. These findings further support the conclusion that differential volume interactions between subunits are mainly responsible for the pattern of contact map compartmentalization.

Our technique allows to validate polymer approaches using chromatin/chromosome condensation algorithm (Andreev and Edel’man, 1999, Zhang and Wolynes, 2015, Brackley et al., 2016, Pierro et al., 2016, Chiariello et al., 2016). Intuitively it is expected that at large (infinite) times the decondensation approach used here to obtain interphase chromosome structure would lead to the same results as the condensation approach. But at small and intermediate times there should be differences. To examine these expectations we calculated the dynamics of gyration radius for the hypothetical process of chromosome 18 condensation under confined environment, Figure 1b, curve 4. Comparison of curves 1 (and 2,3) vs 4 demonstrates that gyration radii asymptotically converge at large time, while at small times they differ significantly and depend on details of mitotic decondensation. Contact maps calculated by the condensation approach (Supplementary Figure 3) are well correlated with our mitotic decondensation (or simply decondensation) approach predictions at the large time (500,000 MC steps). However, at 5,000 - 50,000 MC steps there are visual differences. These data suggest that time required to reach time-independent regime for whole chromosome contact map is different for both approaches, it is longer for condensation than for decondensation pathways. Condensation- and decondensation-based pathways provide the same structure ensemble at large time, whereas at intermediate and small times the differences are significant.

To examine relationships between contact map compartmentalization and spatial compartmentalization of chromosome we design several new proximity functions, T_kn_(x), T(x), F_pq_(x), T_long_(x) and relative proximity functions, f_kn_(x) and g_pq_(x) (see Materials and methods).

T_kn_(x) is a frequency for pairs of subunits of type *k* and type *n* to be separated by 3D distance x, x+dx.

T(x) is a frequency for pairs of subunits of any type to be separated by 3D distance x, x+dx.

F_pq_(x) is a frequency for pairs of subunits from chromosome compartment *p* and compartment *q* to be separated by 3D distance x, x+dx.

T_long_(x) is a frequency for pairs of long-distant subunits (more than 30 Mbp) of any type to be separated by 3D distance x, x+dx.

Functions f_kn_(x) and g_pq_(x) are relative differences, see below in the text.

T_kn_(x) at large distances (˜1 µm) is similar for types 1-1, 2-2, 1-2 (Supplementary Figure 4a) and distinct at small distance corresponding to the range of contact (250-420 nm). To compare T_kn_(x) with T(x) (Supplementary Figure 4b), relative difference function f_kn_(x)=(T_kn_(x)-T(x))/T(x) is calculated. f_kn_(x) is mainly around zero at large distances (Figure 2a). At small distances pairs of the same type are frequent compared to all pairs irrespective of type (Figure 2a, see positive left tail of black and red curves). On the contrary, pairs of different type are rare compared to all pairs (Figure 2a, negative left tail of green curve). These results indicate that subunits of the same type occur in proximity more frequently than those of different types.

To explore how frequently distant compartments of chromosome 18 proximate, we determine distance distribution between pairs of subunits belonging to very distant compartments *p* and *q*, F_pq_(x) (see Materials and methods) and compare with T_long_(x). Compartment numbers p=1, 2, 3, q=19 in Figure 2d; compartments 1 and 19 are shown also in Supplementary Figure 5a,b for artificially decondensed and condensed chromosome. They are positioned on the opposite ends of the chromosome (see the ideogram in Figure 2d) and separated by ˜60-85 Mb. Figure 2b-d indicates that for pairs of compartments 1-19 and 3-19, corresponding to dark regions/increased contact frequency on the map, relative difference g_pq_(x)=(F_pq_(x)-T_long_(x))/T_long_(x) is positive for low distances. For the pair of compartments 2-19, which corresponds to the light region on the map, g_pq_(x) is negative for low distances. That is, these subunits are in proximity less frequently than average over the chromosome. Increased/decreased contact frequencies for distant loci and compartments are the direct consequence of their increased/decreased spatial proximity. Striping of dark/light zones on the map is equivalent to sequence of maximum/minimum on the relative spatial proximity curve (Figure 2c). These results show that distant compartments of the same type proximate with higher frequency than average over the chromosome. Thus the chess-like pattern of contacts reflects spatial compartmentalization of chromosome and vice-versa, but differential volume interactions underlie both effects.

## Discussion

Modeling study of mitosis-G1 transition reveals two-stage dynamics, fast decondensation followed by slow structural rearrangements in G1 accompanied by weak subcondensation. Examination of model dependence of this conclusion reveals that in other scenarios of mitotic decondensation, differing in initial mitotic conformations, the growth of gyration radius is modest but there is subsequent slow decline, as previously. The explanation is simply based on the ratio of the size of the mitotic chromosome and the confinement well.

The next conclusion arising from mitotis-G1 modeling is that the contact map is established soon after decondensation. The maps at early vs late times in G1 are highly correlated and look similar. This conclusion is in accord with the experimental finding that Hi-C interaction maps during interphase (early G1, mid G1, and S-phase) display similar patterns of interactions (Naumova et al., 2013). Some differences of contact maps between early, intermediate and late G1 may be related in the model to further structure equilibration. One cannot exclude an alternative mechanism of early contact map formation, kinetically favorable fast pathway. One can assume some processes like attachment to nuclear matrix *etc.* to be involved. This scenario is omitted in present work.

Our polymer approach to contact map prediction simulates mitosis-G1 transition as a biologically realistic pathway of establishment of interphase chromosome organization and contact maps during G1 phase of cell cycle. It differs from other polymer approaches based on the condensation pathway, irrespective of molecular or coarse-grained algorithms used (Andreev and Edel’man, 1999, Zhang and Wolynes, 2015, Brackley et al., 2016, Pierro et al., 2016, Chiariello et al., 2016). The biological sense of using condensation modeling approaches (based on maximum entropy principle and analogy with protein folding (Zhang and Wolynes, 2015, Pierro et al., 2016), proteins binding to chromatin (Brackley et al., 2016, Chiariello et al., 2016)) is that large scale chromatin/ chromosome condensation is in essence driving force of interphase chromosome organization. That is in contrast with common knowledge that mitotic decondensation in cell cycle results in formation of interphase chromosomes organization. There are approaches that take mitotic decondensation as a starting point for chromosome simulations (Münkel and Langowski, 1998, Rosa and Everaers, 2008, Fritsch and Langowski, 2011). However, these studies do not link the decondensation algorithm with the problem of establishment of interphase contact maps. This problem is solved by the present modeling technique.

The chromosome interactome is not a contact map for single or unique structure. It is calculated as a sum of individual contact maps in cell population. This sum reflects different conformations at a given moment with the same set of potentials in each cell in population. Since thermal motion of polymer subunits during M-G1 decondensation and in interphase is directly captured by our algorithm, a chromosome undergoes thermal fluctuations of conformations in time in each cell. This reveals that ensemble of chromosome conformations arises from stochastic trajectories of chromosomes during mitotic decondensation and thermal fluctuations at any time.

The key Hi-C observations (Lieberman-Aiden et al., 2009, Dixon et al., 2012, Rao et al., 2014), the chess-like contact pattern seen as alternating megabase-sized regions of high vs low intensities on the contact map and spatial compartmentalization of chromosomal subunits, are quantitatively reproduced by mitotic decondensation modeling technique. These observations are explained in the framework of heteropolymer (block-copolymer) chromosome model by differential volume interactions and condensed globular state of a chromosome. Differential volume interactions in the coarse-grained heteropolymer mean that for each type of subunits there is its own pairwise interaction potential. On molecular scale differential volume interactions mean the interactions between certain types of proteins with corresponding types of chromosome subunits but not with others.

Examination of a role of linear memory (“broken chain” calculations) and the external field shows that the specific pattern of contacts remains approximately the same as for “exact” nonbroken polymer chain in the external field, though contact map intensity is different. On the other hand, a homopolymer globule does not reproduce the complex pattern of contacts observed experimentally. Finally, a heteropolymer globular chromosome, placed in decondensation environment, does not show any specific pattern of contacts on the map. These findings further confirm the conclusion that contact and spatial compartmentalization of chromosome is reproduced only if differential volume interactions and globular state of a chromosome exist. This conclusion avoids many biological and biochemical details of chromatin-proteins interactions and arises mainly from basic principles of polymer physics.

A specific parameter introduced here is an external field, see also (Münkel and Langowski, 1998). The need to introduce an external field confining each chromosome in a mammalian nucleus may be motivated by the following arguments. Nuclear organization simulations (Eidelman et al., 2012) demonstrate that all chromosomes in a human interphase nucleus are globular and densely packed, which is consistent with experimental data (Bolzer et al., 2005). Each chromosome is constricted by neighboring chromosomes and/or nuclear shell. This fact can be taken into account as an external field with a shape of a well with rigid or semi-rigid. The Hi-C data description for individual chromosomes without an external field may result in biases in estimating parameters of chromosomes. An external field may reflect a complementary factor shaping chromosomal architecture. One can hypothesize the existence of protein network or structure(s) surrounding each chromosome and playing the role of chromosomal case or external skeleton. This predicts that extraction of chromosomal histone proteins would not disturb a shape of chromosome territory. This idea is not the identical to internal nuclear matrix proposals (Ma et al., 1999). Chromosomal case might provide an additional mechanism of keeping integrity of chromosome territories irrespective to interchromosomal contacts.

One has to note that though our modeling scenario of mitotic-G1 transition is oversimplified from biological stand point, we expect that general conclusions to be model independent. The simulations reproduce the main experimental findings of contact maps (map persistence during G1, specific pattern of contacts).

In conclusion, we present the method demonstrating for the first time how 3D conformation ensemble and interactome for interphase chromosomes are established in the course of mitotis-G1 transition. We identify the physical mechanism of interphase chromosome structure formation which explains main features of Hi-C contact maps for mouse chromosome 18. This mechanism incorporates differential interactions between megabase domains and results in globular state of the interphase chromosome. The mechanism reproduces a chess-like pattern and early G1 establishment of long-range contacts as well as spatial compartmentalization of a chromosome under different conditions. Complementary to volume interactions, a hypothetical chromosomal case may be involved as an important contributor to the early map formation and chromosome integrity.

## Materials and methods

### Chromosome structure and dynamics modeling

Mouse chromosome 18 is represented as a non-lattice coarse-grained chain of 91 semi-rigid spherical 1 Mbp elements corresponding to megabase sized chromosomal subunits, superdomains (SDs) (Eidelman and Andreev, 2002), topological domains (TDs) (Dixon et al., 2012) or topologically associated domains (TADs) (Nora et al., 2012). The first approach, where establishment of the interphase chromosome was modeled in the course of mitotic chromosome decondensation, was introduced (Münkel and Langowski, 1998). Here we link the mitotic decondensation algorithm (Andreev et al., 2006, Eidelman et al., 2006) with different subunit types interactions (block-copolymer composition of the chain) and confining field. This allows to approach the problem of contact map predictions.

The energy of interaction between elements type *i* and *j* through a Lennard-Jones potential is expressed as

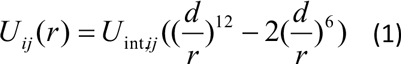

where *r* is distance between subunit centers, *d* is subunit diameter, *U_int, ij_* is energy scale for pair of subunits type *i* and type *j*. Since the chain is a heteropolymer, different types of elements (type 1, type 2, *etc*.) have their own pairwise interaction potentials. For simplicity, *d*=350 nm for all elements. External confining field effectively takes into account interchromosomal interactions. The shape of the field potential is a well with zero potential inside and semi-permeable walls:

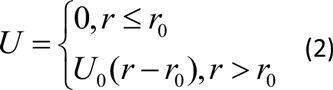

where *r* is distance between centers of the subunit and the chromosome, *r*_0_ is the radius of the well. This type of potential was used for chromosomal aberration modeling (Kreth et al., 2007). The establishment of an interphase chromosome organization is simulated by a gradual equilibration of the chain from different initial conformations. The computer experiments are carried out to explore two scenarios: (a) decondensation from an ultra-compact rodlike structure representing a mitotic-like chromosome; three examples of this scenario are considered by variation of distance between neighboring subunits from 5 to 40 nm to reflect different degree of compactness of mitotic chromosomes; (b) condensation from a real coil. A Metropolis method (Metropolis et al., 1953) with dynamic variant of Monte Carlo (MC) technique is performed, at each MC step positions of elements are randomly changed (Andreev et al., 2006, Eidelman et al., 2006). The simulations were performed for times up to 10 million Monte-Carlo steps at which the macroscopic parameter, chain gyration radius, ceases to change. At different times all macroscopic and microscopic characteristics for individual chromosomes and statistical ensembles (500 structures for each calculation) are obtained.

### Spatial correlation functions

For each pair of subunits (*i*,*j*), placed at positions *i* and *j* on the chain, the space correlation function *t_ij_*(*x*) is introduced as the distribution of 3D distance *x* between subunit centers. Next, the following functions are introduced:

*T*(*x*) is a frequency for pairs of subunits of any type to be separated by 3D distance x, x+dx:

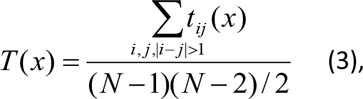

where *N*=91 is the number of subunits in mouse chromosome 18, i.e. *T*(*x*) is *t_ij_*(*x*) averaged over all pairs of subunits except neighbors.

*T_kn_*(*x*) is a frequency for pairs of subunits of type *k* and type *n* to be separated by 3D distance x, x+dx:

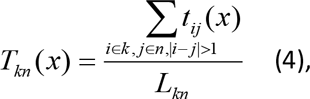

where *L_kn_* is the total number of pairs of subunits type *k* and *n*, i.e. *T_kn_*(*x*) is a pairwise distribution function *t_ij_*(*x*) averaged over positions *i* and *j* on the chain given that they are types *k* and *n*.

*T_long_*(*x*) is a frequency for pairs of subunits with large genomic separation, more than 30 Mbp, of any type to be separated by 3D distance x, x+dx. *T_long_*(*x*) is obtained by averaging of *t_ij_*(*x*) over all pairs of subunits with |*i-j*|>30:

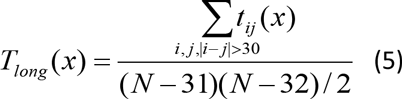

*F_pq_*(*x*) is a frequency for pairs of subunits from chromosome compartment *p* and compartment *q* to be separated by 3D distance x, x+dx. *F_pq_*(*x*) is calculated by averaging of *t_ij_*(*x*) over all pairs of subunits where subunit *i* belongs to compartment *p* and subunit *j* belongs to compartment *q*:

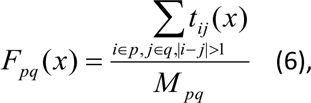

where *M_pq_* is the total number of pairs of subunits from compartments *p* and *q*.

For comparison between different distributions the relative difference functions *f_kn_*(*x*) and *g_pq_*(*x*) are introduced:

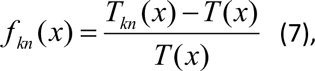

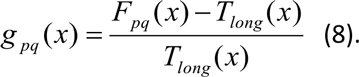

### Contact map calculation

Two elements are deemed contacting if they are in spatial proximity: distance between subunit centers lower than 1.2 diameter (which means 420 nm for subunit diameter 350 nm). In each computer experiment the ensemble of 500 conformations is generated. Total number of contacts between each subunit pair (*i, j*) where |*i-j*|>1, is calculated, i.e. contacts between neighboring subunits are not counted. These numbers of contacts form a table A[91][91]. Then the table is normalized by the total number of contacts to allow for the direct comparison with the experimental data. The experimentally observed contact map or matrix for chromosome 18 is derived from the data set for Hi-C read counts corrected for experimental biases (Rao et al., 2014). The observed contact map is built with 1 Mbp resolution and normalized in the same way as in computer experiments by the total number of counts. No dividing each entry in the contact matrix by the genome wide (or chromosome wide) average contact number of contacts at that genomic distance is used.

### Parameter estimates

The number of element types, the positioning of different types along the chain and the parameters of potentials are determined by an iterative algorithm as follows. 1. For a given set of parameters the simulation is performed, the Pearson correlation between the resulting map and the experimental one is calculated. 2. For element pairs with the highest difference with the experimental data we change either the type or constant for potentials corresponding to their type. Steps 1 and 2 are repeated until the correlation coefficient converges to the maximum. If the correlation coefficient does not converge we increase the number of element types and return to step 1. The initial number of types is 3, one for the near-centromeric 3 Mbp sized region (this is not changed) and two for the other chromosome part. The algorithm proves to converge for 4 types of elements. Examination of type 1 subunits according to their epigenetic signature reveals that they are associated with active chromatin: chromosome 18 regions corresponding to type 1 subunits are enriched with H3K4m1 (histone mark of active chromatin) according to ENCODE data base (ENCODE, 2012). Inspection of type 2-3 subunits indicates depletion of H3K4m1. Type 4 is nearcentromeric chromatin.

## Funding statement

Russian Foundation for Basic Research no. 14-01-00825 to Sergey Andreev

## Funding information

This paper was supported by the following grants:

Russian Foundation for Basic Research no. 14-01-00825 to Sergey Andreev

## Competing interests

The authors declare that no competing interests exist.

